# Direct imaging of neural activity reveals neural circuits via spatiotemporal activation mapping

**DOI:** 10.1101/2024.07.31.606112

**Authors:** Jae-Youn Keum, Phan Tan Toi, Semi Park, Heejung Chun, Jang-Yeon Park

## Abstract

Two years ago, our group reported direct imaging of neuronal activity (DIANA), a functional magnetic resonance imaging (fMRI) technique that directly detects neuronal activity at high spatiotemporal resolution. In this study, we successfully reproduced the DIANA response in medetomidine-anesthetized mice using forelimb electrical stimulation at 11.7 T. More importantly, we showed that multiple neural circuits can be effectively revealed by DIANA fMRI through spatiotemporal activation mapping. The spatiotemporal activation mapping proposed here utilizes the temporal information of the DIANA response, that is, the time when the DIANA response reaches its peak, which is a unique feature that distinguishes it from the activation mapping method used in existing fMRI. Based on DIANA activation areas, we identified several neural circuits involved in forelimb sensory processing in the somatosensory network, which includes multiple brain regions: ventral posterolateral nucleus of the thalamus (VPL), posteromedial thalamic nucleus (POm), forelimb primary somatosensory cortex (S1FL), secondary somatosensory cortex (S2), primary motor cortex (M1), and secondary motor cortex (M2). Additionally, we also identified a pain-related neural circuit involving brain regions of the anterior cingulate cortex (ACC) and mediodorsal nucleus (MD). Interestingly, the spatiotemporal activation mapping also allowed us to identify subregions with different DIANA response times within the same functional region (e.g., VPL, POm, S1FL, and S2). Our study highlights the potential of DIANA fMRI to advance our understanding of sensory information processing throughout the brain and to provide insight into the spatiotemporal dynamics of brain networks at the level of neural circuits.

---

Brain scientists have sought to understand how the brain works in relation to sensation, perception, and action, particularly by attempting to understand the functional organization of the brain, including the dynamic and hierarchical functional connectivity of brain networks (*1–4*). In this regard, since large-scale brain networks consist of multiple neural circuits that are interconnected with each other and perform specific functions, identification of neural circuits, including the causal and temporal ordering of neuronal activity within the circuit, is fundamentally important for implementing realistic dynamic brain network modeling.

To achieve these goals noninvasively *in vivo*, several neuroimaging methods, such as functional magnetic resonance imaging (fMRI), electroencephalography (EEG), and magnetoencephalography (MEG), have been developed. Among them, fMRI meets an important prerequisite for identifying neural circuits by providing good spatial resolution (≤ ∼mm) across large brain areas, including subcortical regions. In particular, blood-oxygenation-level-dependent (BOLD) fMRI has been widely used to investigate brain’s functional organization *in vivo* through neurovascular coupling (*5–7*). Although BOLD fMRI has inherent limitations, such as relatively low temporal resolution due to hemodynamic responses as well as indirect information about neural activity (*8*, *9*), some studies have addressed the temporal order of neural activity in neural circuits using early BOLD responses. For example, several studies using rat forepaw stimulation have reported that early BOLD responses were observed in cortical layer 4 (L4) of the primary somatosensory cortex (S1) (*10–12*). Another recent study by Jung et al. at 15.2 T showed that the sequence of early BOLD responses or cerebral blood volume (CBV) to somatosensory stimulation was consistent with sequential neural information flow in known somatosensory networks, including the ventral posterolateral nucleus (VPL), posterior complex of the thalamic nucleus (PO), S1, secondary somatosensory cortex (S2), and primary motor cortex (M1) (*13*, *14*).

Two years ago, our group reported a novel fMRI method called Direct Imaging of Neuronal Activity (DIANA), which allows direct detection of neural activity on millisecond timescales using a two-dimensional (2D) gradient-echo-based line-scan MR imaging strategy (*15*). DIANA fMRI with 5 ms temporal resolution captured neuronal activity and its propagation in thalamocortical pathways of the somatosensory network during whisker-pad electrical stimulation in anesthetized mice at 9.4 T. The first DIANA response was observed sequentially in the thalamus and primary somatosensory barrel field (S1BF) at ∼10 ms and ∼25 ms, respectively, consistent with previous electrophysiological studies (*16–18*).

Despite the proof-of-concept nature in this first report, DIANA fMRI has shown significant potential to explore dynamic brain networks, including neural information flow, due to its direct correlation with neural activity at millisecond temporal resolution. However, in this first study, DIANA activation was simply identified based on the average of neuroanatomically known functional region of interests (ROIs) such as the thalamus and S1BF. This type of ROI averaging analysis tends to reduce signal sensitivity and specificity because not all neurons in each functional area respond to sensory stimulation every time, and this situation can be further exacerbated when signal sensitivity is relatively low, or when, as in humans, the size of the functional ROI of the brain is much larger and the area is not absolutely defined. Therefore, to make full use of DIANA fMRI when exploring the spatiotemporal dynamics of brain neural networks, it is important to effectively identify DIANA-activated regions in the brain.

In this study, we first proposed an effective data analysis of DIANA fMRI that utilizes temporal information of the DIANA response, which has not been considered in conventional BOLD fMRI, thereby enabling high spatiotemporal resolution activation mapping. Using this new analysis, we were able to identify multiple mouse forelimb sensory circuits involved in the somatosensory network, including both feedback and feedforward pathways (*13*, *14*, *19–28*). For this study, we acquired DIANA fMRI data in medetomidine-anesthetized mice using forelimb electrical stimulation on an 11.7 T animal scanner. In addition to revealing neural circuits using a new analysis method, this study has another important meaning in that it reproduces DIANA fMRI using a different type of sensory stimulation (forelimb vs. whisker pad), anesthetic agent (medetomidine vs. ketamine/xylazine), and magnetic field strength (11.7 T vs. 9.4 T) than the first report of DIANA fMRI.

## Results

### BOLD responses to forelimb stimulation

To confirm the location of main areas of the forelimb sensory circuit, BOLD-fMRI was first performed as a reference on an 11.7 T animal scanner (BioSpec, Bruker) using six adult C57BL/6J medetomidine-anesthetized mice (Fig. 1A). 2D gradient-echo echo planar imaging (EPI) with repetition time (TR) of 1 s and echo time (TE) of 15 ms was used for BOLD fMRI. Electrical stimulation (strength, 0.5 mA; duration, 2 ms; frequency, 10 Hz) consisting of 5 blocks of 20s-on/20s-off cycles were delivered to the right forelimb (Fig. 1B). Because the sensory thalamic nuclei and forelimb primary somatosensory cortex (S1FL) are located in slightly different coronal planes (*29*), ten 0.5 mm thick coronal slices set to include both the thalamus and S1FL were first acquired. As a result, BOLD activation of the thalamus and S1FL was observed in different coronal planes (Fig. 1C for a representative mouse). After examining which coronal slices the thalamus and S1FL responded to, a single 0.8 mm thick oblique slice was acquired using the same electrical forelimb stimulation to observe BOLD responses in the thalamus and S1FL simultaneously (Fig. 1D). BOLD activation maps showed highly consistent activation within the contralateral thalamus and contralateral S1FL across the mice except for the thalamus in mouse #5 (Fig. 1E). Since the oblique slice of mouse #5 appeared to be improperly setup to not include the thalamus, mouse #5 was excluded from further analysis. To confirm the presence of neuronal activity in the thalamus and S1FL, BOLD fMRI using this oblique slice was performed before and after DIANA fMRI. To evaluate the BOLD response in the time course, circular ROIs were selected from the BOLD activation maps in the contralateral thalamus and contralateral S1FL (Fig. 1F). BOLD responses of ∼0.5% were observed in the thalamus and S1FL during the 20 s stimulation-on period (Fig. 1G, *n* = 5 mice), although BOLD responses in the thalamus were weaker than S1FL as reported in previous studies (*13*, *30*).

**Fig. 1.**
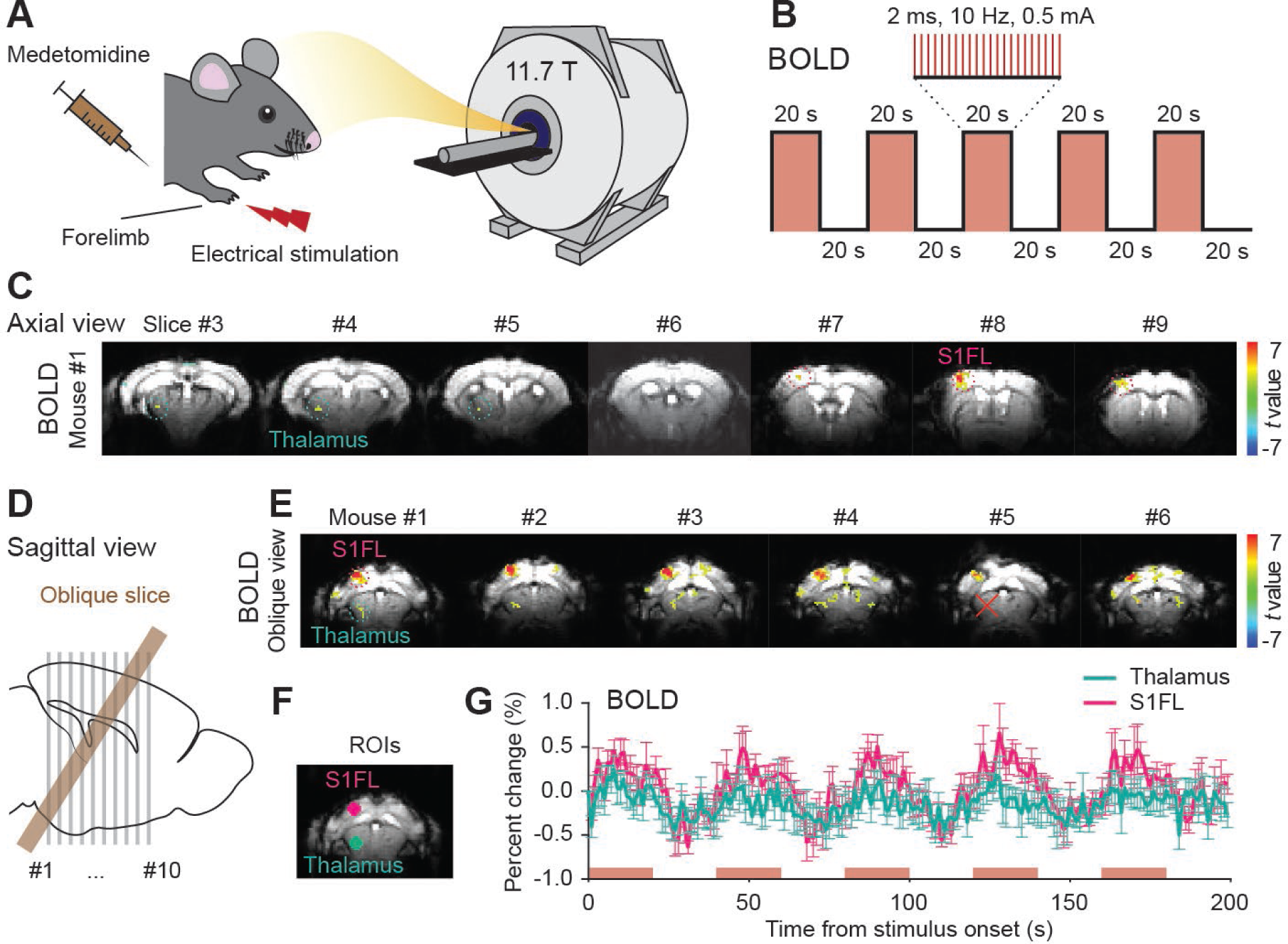
BOLD fMRI in medetomidine-anesthetized mice at 11.7 T. **(A to B)** Schematics of the BOLD fMRI experiment (A) and stimulation paradigm (B) to capture BOLD responses to electrical right forelimb stimulation. **(C)** BOLD responses in coronal slices of a representative mouse (mouse #1). **(D)** Location of the set oblique slice in sagittal plane. **(E)** BOLD responses from 6 mice in the oblique slice. **(F)** Circular ROIs selected for the thalamus (cyan) and S1FL (magenta). **(G)** BOLD time courses extracted from the ROIs of the thalamus and S1FL (*n* = 5 mice). Red box areas indicate the 20s-on state of electrical forelimb stimulation (G). All data are mean ± SEM.

### DIANA responses to forelimb stimulation

To investigate DIANA responses in the mice, the 2D gradient-echo-based line-scan imaging sequence used by Toi et al. (*15*) was implemented on the same 11.7 T animal scanner. 40 trials per mouse were acquired and used for analysis. Sufficient dummy scans (8 s, 1600 TRs) were performed before the main scan to achieve steady-state magnetization (*31*). Both RF spoiling (*32–34*) and gradient spoiling (*35*) were also used to suppress the effects of residual transverse magnetization. Maintaining the same mouse position as the previous BOLD-fMRI experiment, electrical stimulation (strength, 0.5 mA; duration, 1 ms; frequency, 5 Hz) was delivered to the right forelimb to achieve event-synchronized line-scan acquisition (Fig. 2A). The stimulus was applied repeatedly with an interstimulus interval of 200 ms, consisting of 50 ms pre-stimulation, 1 ms stimulation, and 149 ms post-stimulation (Fig. 2B). Stimuli were applied 64 times corresponding to the number of phase-encoding lines. The same oblique slice used in the previous BOLD fMRI experiments was acquired with scan parameters as follows: TR = 5 ms (temporal resolution), TE = 2 ms, flip angle = 4°, and spatial resolution = 0.2×0.2×0.8 mm^3^ (Fig. 2C). To perform group analysis on 5 mice, each DIANA image for mice #2 to #6 (except mouse #5 due to no BOLD response in the thalamus) was linearly registered to the DIANA image for mouse #1 and then averaged (Fig. 2D, leftmost).

**Fig. 2.**
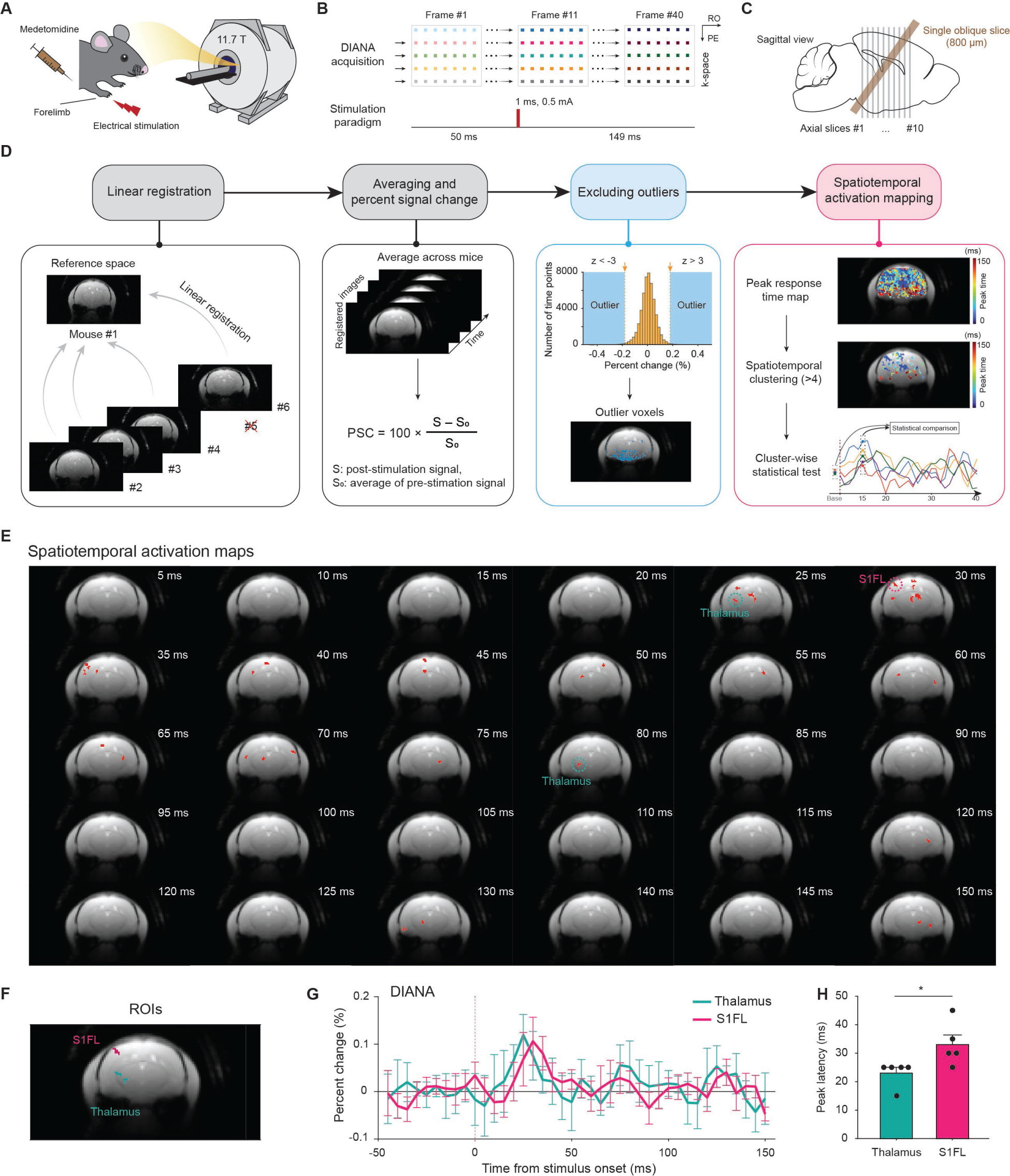
Spatiotemporal activation mapping in DIANA fMRI. **(A to B)** Schematics of the DIANA fMRI experiment (A), 2D line-scan acquisition and stimulation paradigm (B) to capture DIANA responses to electrical right forelimb stimulation in medetomidine-anesthetized mice at 11.7 T. **(C)** Location of the set oblique slice in sagittal plane. **(D)** Flowchart of the spatiotemporal activation mapping process: Linear registration of DIANA images of 4 mice (#2 to #6, excluding #5 due to no BOLD response in the thalamus) to the mouse #1 image as template. Average the data across all mice and represent the time series as percent signal change (PSC). Exclude outlier voxels with z > 3 or z < −3 using a histogram of the number of voxels with respect to PSC. Spatiotemporal activation mapping consisting of generation of a peak response time map, spatiotemporal clustering (cluster size > 4), and cluster-wise testing for statistical significance of DIANA responses. **(E)** Spatiotemporal activation maps of DIANA fMRI with 5 ms temporal resolution after stimulus onset. **(F)** Representative DIANA-activated regions within the thalamus (cyan) and S1FL (magenta). **(G to H)** DIANA time series (G) and mean latencies (H) of DIANA response peaks obtained from the DIANA-activated regions within the thalamus and S1FL (*n* = 5 mice, *: *p* < 0.05, unpaired *t*-test). Vertical dotted lines indicate the onset time of electrical forelimb stimulation (H). All data are mean ± SEM.

For data analysis, we developed a data analysis method (termed “spatiotemporal activation mapping”) to identify DIANA activation regions in an effective and reliable manner by exploiting the temporal information of DIANA responses, which is a key advantage of DIANA fMRI over conventional fMRI (Fig. 2D). In other words, the peak response time map was created by obtaining the times of peak DIANA responses on a voxel-by-voxel basis from the time series of the averaged DIANA image, and clustered voxels above a cluster size of 4 were selected as activation regions. An important rationale for this spatiotemporal activation mapping is the assumption that spatially clustered voxels of a certain size in the peak response time map can be considered a group of functionally identical features sharing the same response time and cannot be generated by chance. As a final step, we statistically compared the DIANA signal at all post-stimulus time points to the pre-stimulus baseline to test the statistical significance of the DIANA response for the mean time series of each cluster. Clusters that survived this statistical test were considered to represent the final spatiotemporal activation areas.

Prior to the spatiotemporal activation mapping, voxels with at least one data point with z score > 3 or < −3 were excluded as outliers, considering the percent signal change (PSC) histogram of all data points in the time series for all voxels. Here, the PSC was defined as a percentage of the post-stimulus measurement relative to the average of the data points in the pre-stimulus period. The outlier voxels were mostly concentrated in the lower part of the brain (Fig. 2D, second from the right), where the temporal signal-to-noise ratio (tSNR), defined as the signal mean divided by signal standard deviation (SD) over the 40 frames, was relatively low because a surface coil was used on top of the brain (Supplementary Fig. S1A). This is because high PSC values corresponding to z score > 3 or < −3 may be due to low average values in the pre-stimulus period in the low tSNR region. The maximum and minimum values of the tSNR map obtained from the average time series of 200 trials (= 5 mice × 40 trials/mouse) were 1469.4 and 104.17, respectively. The mean tSNR value of the outlier voxels was 371.33 ± 104.04.

As a result, the spatiotemporal distribution of all DIANA activation areas was shown in Fig. 2E, including the somatosensory network. Among all activation areas, the thalamocortical pathway from the ventral posterolateral nucleus (VPL) of the thalamus to S1FL was first presented in Fig. 2F to H, representatively showing the reproduction of DIANA fMRI at 11.7 T. In response to the electrical forelimb stimulation, statistically significant DIANA signals were observed in the VPL and S1FL compared to a control region in the muscle (*p* < 0.0001 for the thalamus and *p* < 0.001 for S1FL, *n* = 5 mice) (Supplementary Fig. S1). Peak DIANA responses occurred in the order of the VPL and S1FL (Fig. 2G) with latencies of 23.0 ± 2.00 ms and 33.0 ± 3.39 ms, respectively (Fig. 2H, *n* = 5 mice), which was in good agreement with previous electrophysiological studies using mouse forelimb stimulation (*21*, *36–42*). It is interesting to note that there was a 5 - 15 ms delay in the DIANA response to forelimb stimulation compared to whisker pad stimulation reported in (*15*), probably due to the longer pathway of the forelimb sensory circuit.

### Forelimb sensory pathways revealed by DIANA fMRI

Next, we investigated whether previously reported forelimb sensory circuits (*13*, *14*, *19*–*28*), including feedforward and feedback pathways, could be identified from the spatiotemporal activation maps in Fig. 2E. Referring to the Allen Mouse Brain Atlas (*29*) (Fig. 3A), we investigated several DIANA activation regions associated with the somatosensory network: VPL, posterior medial thalamic nucleus (POm), S1FL, S2, M1, and secondary motor cortex (M2) (Fig. 3B). To find forelimb sensory pathways, we used DIANA response time for each activation region, along with known causal relationships between somatosensory network regions (*13*, *14*, *19–28*). All peak responses corresponding to DIANA response times of these activation areas were statistically significant relative to the mean of the pre-stimulation period, and statistical significance was indicated on top of each peak in the time series (Fig. 3C to H) (*: *p* < 0.05, **: *p* < 0.01 for one-tailed Wilcoxon signed-rank test for peak response of MD and one-tailed Welch’s *t*-test for all peak responses except for MD, See Supplementary Fig. S2 for more details).

**Fig. 3.**
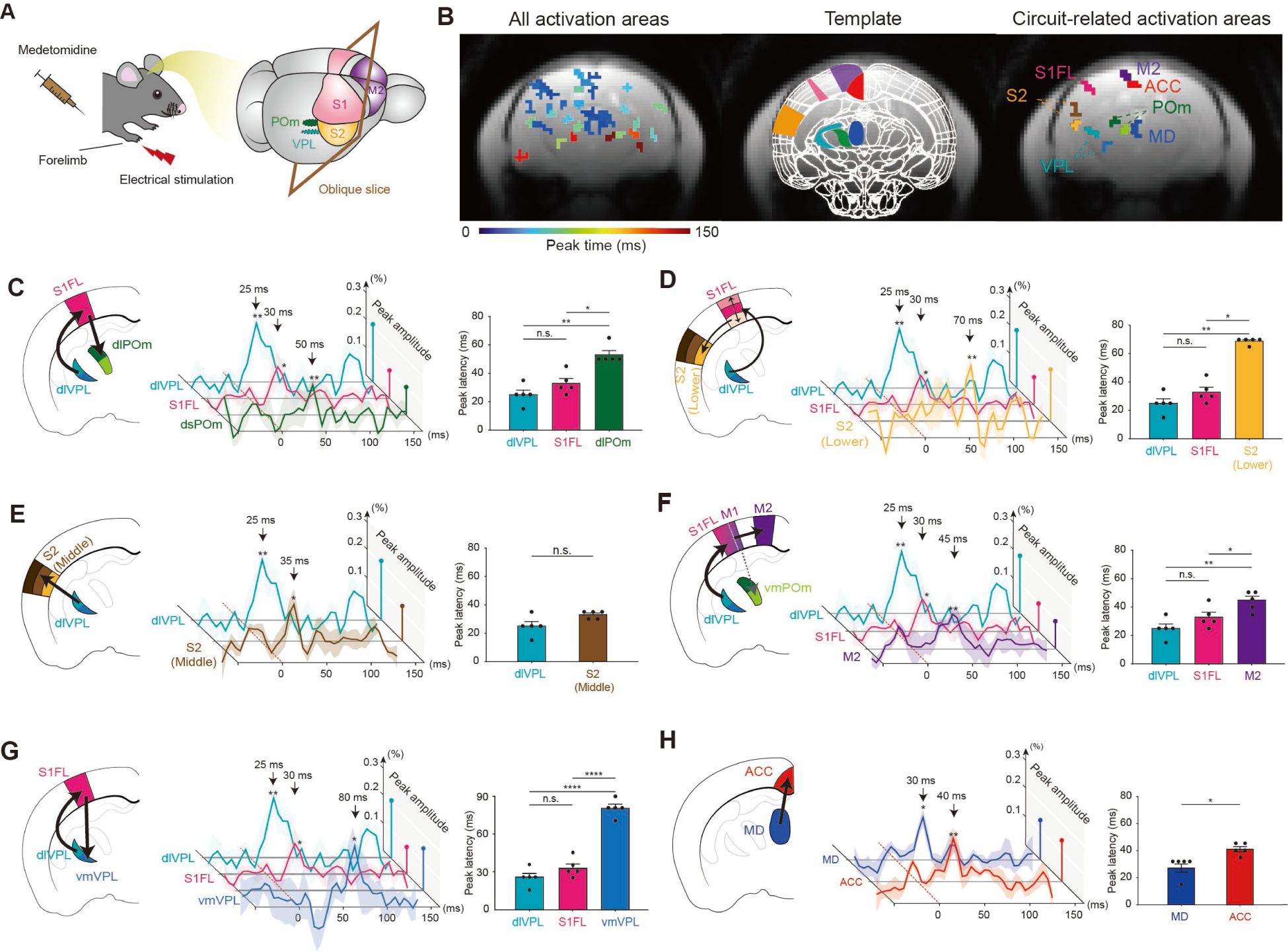
DIANA fMRI reveals multiple neural circuits in the somatosensory network in response to mouse forelimb stimulation. **(A)** Schematics of the DIANA fMRI experiment (left) and targeted oblique slice covering VPL, POm, somatosensory cortex, and motor cortex (right). **(B)** All activation areas obtained through spatiotemporal activation mapping (left), where colors represent peak response times. Allen Mouse Brain Atlas template matched to the oblique DIANA image (middle) and activation areas associated with the neural circuits presented below (right), where the colors represent different activation areas and are different from the colors used for peak response times (left). **(C) Circuit I** (VPL ➔ S1FL ➔ POm): Illustration of TC and CT projections from the dorsolateral VPL (dlVPL, cyan) through S1FL (magenta) to the dorsolateral POm (dlPOm, green) (left). DIANA time series (middle) and mean latencies (right) of peak DIANA responses in **Circuit I** are also shown (*n* = 5 mice). **(D) Circuit II** (VPL ➔ S1FL ➔ S2): Illustration of TC and CC projections from the dlVPL through S1FL to the lower S2 (yellow) (left). DIANA time series (middle) and mean latencies (right) of peak DIANA responses in **Circuit II** are also shown (*n* = 5 mice). **(E) Circuit III** (VPL ➔ S2): Illustration of TC projection from the dlVPL to the middle S2 (brown) (left). DIANA time series (middle) and mean latencies (right) of peak DIANA responses in **Circuit III** (*n* = 5 mice). **(F) Circuit IV** (VPL ➔ S1FL ➔ M1* ➔ M2): Illustration of TC and CC projections from the dlVPL through S1FL/M1* to the M2 (purple) (left). DIANA time series (middle) and mean latencies (right) of peak DIANA responses in **Circuit IV** are also shown (*n* = 5 mice). **(G) Circuit V** (VPL ➔ S1FL ➔ VPL): Illustration of CT and TC projections from the dlVPL through S1FL back to the ventromedial VPL (vmVPL, blue), as a recurrent loop (left). DIANA time series (middle) and mean latencies (right) of peak DIANA responses in **Circuit V** are also shown (*n* = 5 mice). **(H) Circuit VI** (MD ➔ ACC): Illustration of TC projection from the MD (navy) to the ACC (red). DIANA time series (middle) and mean latencies (right) of peak DIANA responses in **Circuit VI** (*n* = 5 mice). In time series plots, the maximum DIANA response amplitude for each ROI was displayed as a percentage change in the right-end plane in projection form. Dotted red lines indicate the onset time of electrical forelimb stimulation. All data are mean ± SEM. *: *p* < 0.05, **: *p* < 0.01, ****: *p* <0.0001, n.s.: *p* > 0.05 for Kruskal-Wallis ANOVA with LSD *post hoc* test (bar graphs in C and D), one-way ANOVA with Fisher’s LSD *post hoc* test (bar graphs of F and G), one-tailed Wilcoxon signed rank test (bar graphs in E, H, and peak response of MD), and one-tailed Welch’s *t*-test (all peak responses except for MD).

First, we identified the forelimb sensory circuit connecting the VPL to S1FL via thalamocortical (TC) projections (*13*, *14*, *19–22*) and subsequently to the POm, a higher-order thalamic nucleus in the somatosensory system, via a corticothalamic (CT) projection (**Circuit I**: VPL ➔ S1FL ➔ POm, Fig. 3C) (*23–26*). The feedforward TC projection from the dorsolateral VPL (dlVPL) to S1FL and the feedback CT projection to the dorsolateral POm (dlPOm) were sequentially observed, with peak latencies of 25.0 ± 3.16 ms (dlVPL), 33.00 ± 3.39 ms (S1FL), and 53.0 ± 3.00 ms (dlPOm), respectively (*n* = 5 mice). Second, the corticocortical (CC) pathway from S1FL to the S2 was found (*13*, *14*, *27*), with a peak latency of 69.0 ± 1.00 ms in the lower S2, following the TC projection to S1FL (**Circuit II**: VPL ➔ S1FL ➔ S2, Fig. 3D) (*n* = 5 mice). Third, the feedforward TC projection connecting the dlVPL directly to the S2 was also found (**Circuit III**: VPL ➔ S2, Fig. 3E) (*14*, *20*), with a peak latency of 33.0 ± 1.22 ms in the middle S2 (*n* = 5 mice). Fourth, following the feedforward TC projection from the dlVPL to S1FL, the subsequent CC projection to M2 (*43*) was also identified with a peak latency of 44.0 ± 2.92 ms (**Circuit IV**: VPL ➔ S1FL ➔ M1* ➔ M2, Fig. 3F) (*n* = 5 mice). Here, S1FL and M2 activation was clearly observed, but M1 activation was only observed in a region slightly overlapping with S1FL and was not separated from S1FL activation in the peak response time map. Previous studies using voltage-sensitive dye imaging have shown that the contralateral forepaw motor area can overlap by ∼50% with the contralateral forepaw somatosensory area (*44*), suggesting that S1 and M1 activation may occur in nearly overlapping activation areas. Fifth, the cortico-thalamo-cortical (CTC) circuit as a recurrent loop that has another CT projection from the S1FL back to the VPL was also found (**Circuit V**: VPL ➔ S1FL ➔ VPL, Fig. 3G) (*19*, *22*), with a peak latency of 80.0 ± 3.16 ms in the ventromedial VPL (vmVPL) (*n* = 5 mice). Finally, we also found another TC projection involved pain processing, connecting mediodorsal nucleus (MD) of the thalamus to the anterior cingulate cortex (ACC) (**Circuit VI**: MD ➔ ACC, Fig. 3H) (*45*, *46*), with peak latencies of 27.0 ± 3.00 ms (MD) and 41.0 ± 1.87 ms (ACC), respectively (*n* = 5 mice). MD is known to be involved in nociception due to its large number of nociceptive neurons and (*47*, *48*), and ACC is known to be connected to MD and play an important role in pain processing (*49*).

Together, spatiotemporal activation mapping of DIANA responses was able to reveal multiple mouse forelimb sensory circuits *in vivo*, including feedforward TC and CC projections as well as feedback CT projections. The connectivity of all six circuits from **Circuit I** to **VI** has already been shown using neuronal tracers (*13*, *14*, *19*, *20*, *22*, *25*, *27*, *49–51*). In addition, the temporal order of neuronal events in these six circuits have also been validated *in vivo* by using BOLD and CBV fMRI (**Circuit I, II, III, IV\***) (*13*, *14*) and electrophysiological recordings (**Circuit I, III, IV, V, VI**) (*19–21*, *23–25*, *46*).

It is interesting to note that activation areas within the POm and VPL of the thalamus and S1FL and S2 of the somatosensory cortex were observed as separate subregions. For example, the dlPOm of **Circuit I** (VPL ➔ S1FL ➔ POm) showed faster responses (53.0 ± 3.00 ms), whereas the vmPOm showed slower responses (69 ± 4.85 ms), which could be considered a feedback CT projection from M1 to vmPOm in **Circuit IV** (VPL ➔ S1FL ➔ M1* ➔ M2, Supplementary Fig. S3) (*19*, *51*). This is because neural activity propagation would take slightly longer to undergo a CC projection to M1 after S1FL activation compared to **Circuit I**.

For the VPL, the response time of the dlVPL in **Circuit I** to **IV** (25.0 ± 3.16 ms) was consistent with previous electrophysiological studies using forelimb sensory stimulation (*36*, *41*), and thus the vmVPL with slow responses (80.0 ± 3.16 ms) was considered to be involved in **Circuit V** (VPL ➔ S1FL ➔ VPL), a CTC circuit as a recurrent loop (*19*).

For the S1FL, as shown in the spatiotemporal distribution of DIANA activation in Fig. 2E, S1FL activation occurred at the 30 ms point in the time series (actually 33.00 ± 3.39 ms) in the middle part of S1FL, subsequently followed by activation at the 35 ms point in the time series in the outer part of S1FL, i.e., the upper (36.00 ± 3.32 ms) and lower (39.00 ± 2.92 ms). Because the middle part can be considered to correspond to layer 4 (L4), the upper part to layer 2 and 3 (L2/3), and the lower part to layer 5 (L5), this information flow between cortical layers in S1 is in good agreement with previous studies using forelimb or whisker stimulation (*13*, *14*, *27*, *50*).

In addition, S2 activation was separated into a middle region with faster responses (33.0 ± 1.22 ms) and a lower region with slower responses (69.0 ± 1.00 ms). The lower and middle S2 regions were considered to be involved in **Circuit II** and **Circuit III**, respectively, because a single TC projection (VPL ➔ S2) in **Circuit III** was expected to be shorter than a sequential combination of TC (VPL ➔ S1FL) and CC (S1FL ➔ S2) projections in **Circuit II**.

In **Circuit V**, we also observed a significant decrease in the DIANA signal in the vmVPL immediately before a significant signal increase in the dlVPL (Fig. 3G and Fig. 4C). We interpret that the signal decrease in the VPL region may be due to hyperpolarization of neuronal membrane potential, such as occurs in cases of GABAergic inhibition by nearby interneurons. The GABAergic inhibition by local interneurons in the relay nucleus of the thalamus is known to limit the central excitatory region of the thalamus, thereby suppressing irrelevant information and leading to accurate sensory processing (*52–54*). These processes can occur through feedforward or feedback recurrent circuits between the cortex and thalamus and can be the underlying mechanisms of attention and goal-directed behavior (*55*, *56*). In addition, neuronal hyperpolarization in the VPL region is known to trigger activation of hyperpolarization-activated cyclic nucleotide-gated (HCN) channels and the occurrence of rebound spike-burst firing of thalamocortical neurons in the thalamus-cortex recurrent loop (*57–59*).

**Fig. 4.**
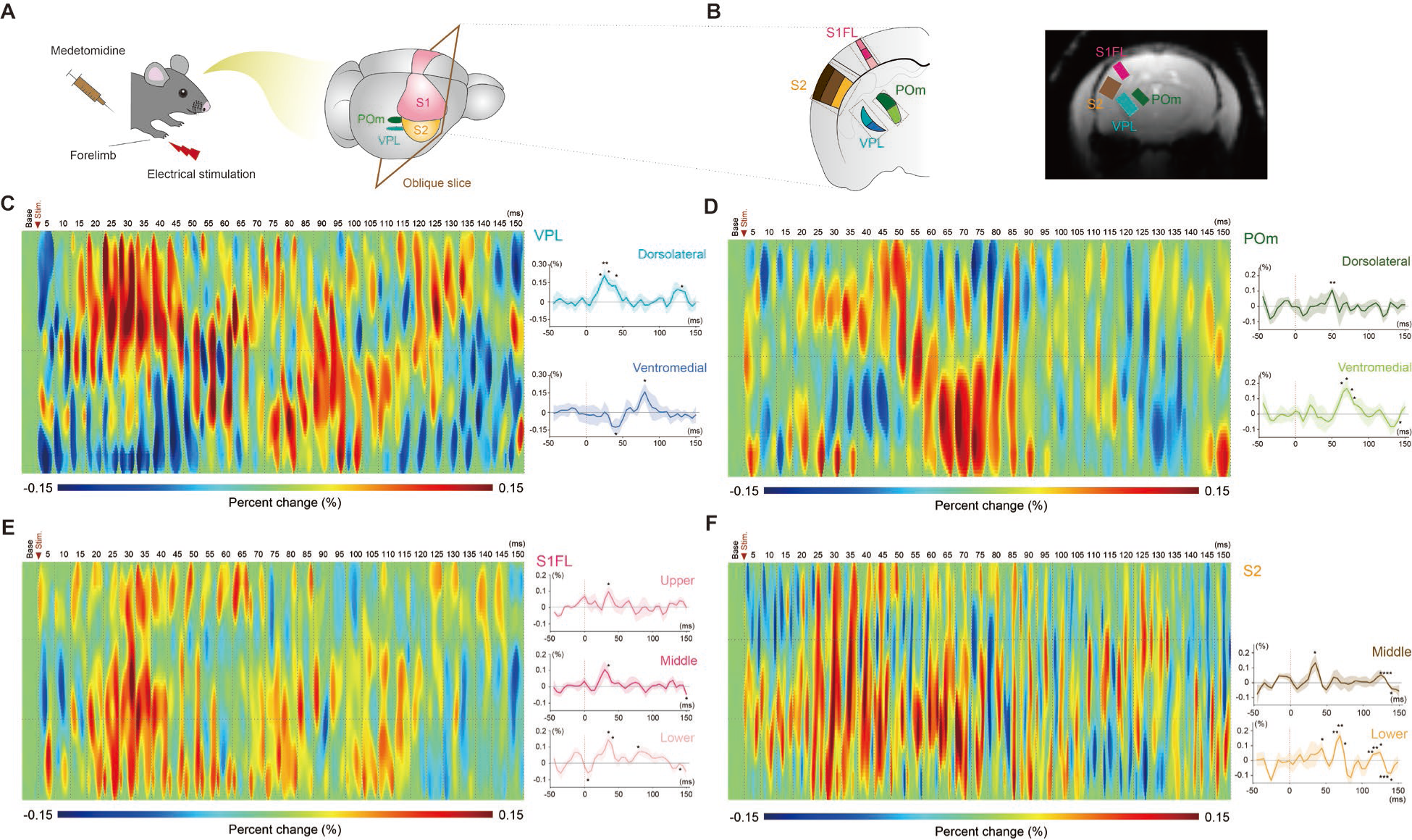
Subregion-specific DIANA responses to mouse forelimb stimulation. **(A)** Schematic of the DIANA fMRI experiment (left) and an oblique slice including the VPL, POm, somatosensory cortex (right). **(B)** Illustration of a brain atlas of the oblique slice (left) and rectangular boxes (gray dotted line) set to include the DIANA-activated region of each ROI (right). The DIANA activation heatmap below is shown in the area within the rectangular box. **(C)** Heatmap of the VPL (left) and time series of two subregions (dorsolateral and ventromedial) of the VPL (right). **(D)** Heatmap of the POm (left) and time series of two subregions (dorsolateral and ventromedial) of the POm (right). **(E)** Heatmap of the S1FL (left) and time series of three subregions (upper, middle, and lower) of the S1FL (right). **(F)** Heatmap of the S2 (left) and time series of two subregions (middle and lower) of the S2 (right). The background of the heatmap represents the average of the pre-stimulation period. The black horizontal dotted lines in the heatmap indicate the tentative boundaries of the subregions. The red vertical dotted lines in the time series indicate the onset time of electrical forelimb stimulation. *: *p* < 0.05, **: *p* < 0.01, ***: *p* < 0.005, ****: *p* <0.0001 for one-tailed Wilcoxon signed-rank test when normality is not followed and one-tailed Welch’s *t*-test when normality is followed.

With respect to **Circuit VI**, MD has been reported to receive nociceptive inputs from the spinal cord (*60*, *61*), the medullary subnucleus reticularis dorsalis (*62*), and the pontine parabrachial nucleus (*63*). This suggests that MD is activated through pathways independent of the VPL. Additionally, a previous study stimulating the MD and recording an excitatory postsynaptic potential (EPSP) in the ACC showed that a ∼10 ms delay was observed in the ACC after MD stimulation (*46*), which is consistent with our findings in **Circuit VI**.

To better represent the spatiotemporal dynamics of functional regions containing separate activation areas, heatmaps were generated by selecting the minimal rectangular ROI containing the activation areas in each of VPL, POm, S1FL, and S2 (Fig. 4). For better heatmap representation, spatial resolution was increased by 10× and additional spatial smoothing was performed. As shown in the time series analysis (Fig. 3), it is visually well represented that peak DIANA responses in both VPL and POm were activated first in the dorsolateral (dl) region and then in the ventromedial (vm) region (Fig. 4, C and D). It is also well visualized that peak DIANA responses in S1FL originated in the middle layer, followed the lower and upper layers (Fig. 4E), while in S2, the middle layer was activated first and then the lower layer (Fig. 4F).

## Discussion

Here, we successfully reproduced DIANA responses using forelimb electrical stimulation in medetomidine-anesthetized mice at 11.7 T, averaging all trials across responded mice (40 trials/mouse, 5 mice). Furthermore, we proposed a new analysis to obtain spatiotemporal activation maps from DIANA fMRI using temporal information of neuronal activation, a concept not previously used in fMRI studies. Through this new analysis, we were able to effectively reveal multiple neural circuits of the somatosensory system involved in forelimb sensory information processing with a temporal resolution of 5 ms. This study highlights the exciting ability of DIANA fMRI to noninvasively and dynamically reveal *in vivo* neural circuits of the brain network in large brain regions with ultrahigh spatiotemporal resolution, in marked contrast to other neuroimaging techniques.

It is particularly interesting to note that spatiotemporal DIANA activation mapping can isolate activated regions with different response times, even within the same neuroanatomical region. For example, after S1FL activation at 30 ms sampling time, mainly in the middle of the S1FL region, propagation of neural activity to the outer layers of S1FL at 35 ms sampling time was observed over the course of the CC and CT pathways (*13*, *14*, *27*, *50*). Higher-order S2 activation was also observed in two separate subareas of S2, with fast responses (33.0 ± 1.22 ms) in the middle part and slow responses (69.0 ± 1.00 ms) in the lower part. Because, in S2, layer 4 (L4) is known to receive input through the TC pathway and layer 2/3 (L2/3), layers 5 (L5) and layer 6 (L6) receive input through the CC pathways (*14*, *27*, *50*), the early DIANA response in the middle S2, assumed to correspond to L4, suggests a TC input, while the later response in the lower S2, where L5 and L6 are assumed to be located, suggests a CC input. Interestingly, the peak response of S2 via the CC pathway (∼0.19 %) was approximately 1.5 times higher than that via the TC pathway (∼0.13%), indicating that S2 is more responsive to CC input delivered from the S1FL than TC input delivered from the VPL (*14*).

In addition to the somatosensory cortical regions of S1FL and S2, the sensory thalamic nuclei of VPL and POm each showed two separate activated subregions with different DIANA response times, respectively. The dlVPL, considered to be involved in the feedforward TC pathway to S1FL, showed early responses (25.0 ± 3.16 ms), while the vmVPL showed later responses (80.0 ± 3.16 ms) through a feedback CT pathway from S1FL (*19*, *51*). POm activation was first observed at 53.0 ± 3.00 ms in the dlPOm, and another activation was later observed at 69 ± 4.85 ms in the vmPOm. The early POm activation in the dlPOm was considered to be due to a feedback CT projection from S1FL (S1FL ➔ POm), and later activation in the vmPOm was considered to be due to another feedback CT projection from M1* (M1* ➔ POm) via an additional CC projection (S1FL ➔ M1*).

It is also worth mentioning that a neural circuit, which can be considered a cortico-striatal (CS) projection connecting the S1FL and the caudoputamen (CP), was also found in the spatiotemporal activation map (VPL ➔ S1FL ➔ CP, Supplementary Fig. S4). This circuit has previously been identified through neural tracing and is known to play a critical role as the initial input from the cortex to the basal ganglia and to influence sensorimotor, cognitive and emotional functions (*64*, *65*). Interestingly, four activated regions within the CP were identified, of which the lateral region with the highest DIANA response showed a peak latency of 59.0 ± 2.45 ms (Supplementary Fig. S4). The lateral CP, especially the ventrolateral region of the medial CP, is known to be involved in the implementation of stereotyped behaviors of the upper limb (*65*). The function and meaning of the remaining three activated regions in CP require further investigation.

Since DIANA fMRI has been proposed to exhibit direct correlation with neuronal activity at millisecond temporal resolution, it appears to have some notable implications that previous fMRI studies did not have to consider in terms of data acquisition and analysis. For example, according to our observations, it seems that averaging more trials in a single mouse may not always ensure higher sensitivity of the DIANA response. This observation appears to contradict the well-known principle that signal averaging improves the signal-to-noise ratio (SNR), usually calculated by dividing the main signal by the standard deviation (SD) of the background noise (*66*, *67*). For example, averaging *N* repeated measurements improves SNR by 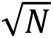 because the sum of *N* measurements increases the main signal by *N* and the SD of the noise by 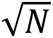. However, this well-known principle is based on two fundamental assumptions, that is, the consistency of the main signal and the randomness of the noise. In terms of the consistency of the main signal, due to the direct correlation of the DIANA response with neuronal activity and its relatively small percent change, the variability of neuronal responses, including neuronal adaptation (*68*, *69*), should be taken more seriously in DIANA-fMRI compared to conventional BOLD-fMRI. In addition, our signal analysis showed that baseline fluctuations in the DIANA signal contained pseudorandom signal components and were suppressed more effectively by averaging data across mice than by averaging the same number of data from a single mouse (Supplementary Fig. S5A). One possible scenario is that if the DIANA signal from each mouse has non-random oscillatory signal components with specific frequency and phase ranges, these signal components could destructively interfere with each other and be effectively suppressed when averaged across mice. (Supplementary Fig. S5B and C). Taken together, in terms of data acquisition, short breaks (e.g., ∼60 - 90 s) are practically recommended every few trials to minimize significant neuronal adaptations. In terms of data analysis, to detect the much smaller amplitude of the DIANA response (e.g., ∼0.1 – 0.2%), it is recommended to average the data across mice (e.g., ≥ ∼5 mice), collecting a sufficient number of trials (e.g., ≥ ∼a total of 200 trials for all mice) such that the tSNR of the averaged data is much higher for DIANA fMRI than for BOLD fMRI. The average tSNR of all voxels without outliers was 626.91 ± 175.30 in this study.

From the perspective of MR physics, SNR increases with increasing magnetic field strength. However, although a higher magnetic field of 11.7 T was used in this study, the percent signal change of the DIANA response was ∼0.1 - 0.2%, which is similar to, but not better than, the previously reported results at 9.4 T. There may be a couple of possible reasons for this. One possible reason could be the use of another anesthetic, namely medetomidine, instead of ketamine, which is known to reduce bottom-up information processing by inhibiting norepinephrine neurons in the locus coeruleus (*70*). A previous study has shown that somatosensory BOLD-fMRI responses under ketamine are stronger than under dexmedetomidine (*71*). Another possible reason is that the mouse brain may be less responsive to forelimb stimulation compared to whisker-pad stimulation. In our experiment performed at a higher magnetic field of 11.7 T, not only DIANA responses, but also BOLD responses were lower (∼0.5%) than responses to whisker pad stimulation at 9.4 T (∼2 - 3%) (*15*). A more fundamental reason may be related to the underlying signal mechanism of DIANA fMRI. As suggested in previous studies (*15*, *72*), the contrast mechanism of DIANA fMRI may be changes in spin-spin relaxation time (T_2_) due to changes in membrane potential that occur during neuronal activation. For DIANA-fMRI using the 2D gradient-echo-based line-scan imaging technique, changes in T_2_ are reflected as changes in apparent T_2_, which is called T * that includes the effects of magnetic field inhomogeneity. As magnetic field strength increases, the T_2_* can be shorter, which may work against the increase in SNR, when using the same echo time at higher magnetic field strengths. Using a shorter echo time can compensate for the effect of T_2_* shortening in terms of signal sensitivity, but at the same time reduces T_2_ contrast, a possible source of DIANA signal changes. Therefore, there may be an optimal magnetic field strength and parameter settings that compromise between T * shortening and T_2_ contrast. This may be one of the reasons why DIANA-fMRI replication was not successful in another recent study using a much higher magnetic field strength of 15.2 T with the same parameter settings (*73*) as the original DIANA report at 9.4 T (*15*).

On the other hand, there are also criticisms that the DIANA signal may be an artifact signal due to nonideal aspects of the pulse sequence. Valerie Doan et al. reported that the delay time assigned to trigger stimulation in the DIANA pulse sequence can disrupt the steady state of the spin system and cause undesired signals that are not related to neuronal activation (*74*). They made the important discovery that even small trigger delays (e.g., 12 μs) in the line-scan pulse sequences of DIANA fMRI can produce unwanted signals, the amplitude of which decreases with decreasing T_2_ of the imaging object. However, if the trigger delay and T_2_ value are not too long, these unwanted signals can be suppressed by sufficiently spoiling the transverse magnetization, i.e. by applying RF spoiling as well as gradient spoiling. For example, when using a water phantom (T_2_ = 1.25 s) at 9.4 T, with a trigger delay of 25 μs (as in our case), a signal change with a peak of 0.083% was observed 15 ms after trigger onset when RF spoiling was turned off, whereas a signal change peak of 0.056% was observed 10 ms after trigger onset when RF spoiling was turned on (Supplementary Fig. S6A and B). However, when using a water phantom with trace amounts of gadolinium (T_2_ = 350 ms) with RF spoiling on, signal changes due to trigger delay were well suppressed (0.00080 ± 0.0066% for all time points, Supplementary Fig. S6C). On the other hand, T_2_ values of mouse or rat brain cortex at 9.4 T and 11.7 T range from 30 to 50 ms (*75–77*), and as T_2_ decreases, the amplitude of signal change due to trigger delay decreases. Therefore, if RF spoiling is turned on and the trigger delay is not that long (e.g. < 30 μs), we can reasonably say that undesired signals due to the trigger delay will not have a significant effect on mouse/rat DIANA fMRI.

There are several points in this study that need to be discussed: First, due to the difficulty in normalizing an oblique slice to the mouse brain atlas, for group analysis of five mice in this study, instead of registering each slice from five mice to the mouse brain atlas template, we linearly registered each slice of the four mice (#2 to #6, excluding #5) to the slice of mouse #1, and then manually found an oblique template that matched the slice of mouse #1 as closely as possible from the Allen Mouse Brain Atlas (Fig. 3B). Considering the convenience of automatically registering images to the brain atlas template, if the study design allows, it seems more desirable to establish single or multiple coronal slices rather than oblique slices to include brain regions relevant to the neural circuits of interest. Second, because it is well known that sensory stimulation elicits more pronounced responses in contralateral somatosensory networks, this study focused on forelimb sensory circuits involving contralateral brain regions. However, through the spatiotemporal activation mapping, DIANA responses were also observed in ipsilateral brain regions such as MD, ACC, M1, CP, reticular nucleus of the thalamus, S2, and ventral posteromedial nucleus (Fig. 2F). Further investigation of these ipsilateral responses in relation to contralateral responses, including callosal projections (*78–80*), would also be an interesting study. Finally, we identified several forelimb sensory circuits (**Circuits I** to **VI**) of the somatosensory network based on known network information, including neuroanatomical and functional connectivity (*13*, *14*, *19–25*, *27*, *46*, *49–51*). However, using spatiotemporal DIANA activation maps to reveal unknown or more complex neural networks in large and distant brain regions may require not only knowledge of known neural networks, including neuroanatomical connectivity, but also dynamic causal models to properly determine the causal relationships in neural networks, including Granger causality (*81*) and transfer entropy (*82*).

In conclusion, we successfully observed DIANA responses in the forelimb sensory circuits of the somatosensory system *in vivo* using electrical forelimb stimulation in anesthetized mice on an 11.7 T animal scanner. Sequential DIANA responses were also identified in forelimb sensory circuits, and the temporal order of DIANA responses was consistent with previously known information flow in the mouse somatosensory network. DIANA fMRI is expected to make a significant contribution to revealing the brain’s neural circuits and thereby creating a realistic dynamic brain network model with high spatiotemporal resolution.

## Acknowledgments

We thank J. Valette and C. Baligand (MIRCen, CEA) for arranging the experiments at 11.7 T and for valuable scientific discussion; S. Malaquin, C. Hery, and E. Mougel (MIRCen, CEA) for preparing and helping experiments. We also thank M. Lowe, W. Shin, A. Nemani, K. Sakaie (Imaging Institute, Cleveland Clinic) for insightful scientific discussion.

## Funding

J.-Y.K., P.T.T., S.P., and J.-Y.P. acknowledge financial support by the National Research Foundation of Korea (NRF) grant funded by the Korea government (MSIT): NRF-2019M3C7A1031993 and 2023R1A2C3007075.

## Author contributions

Conceptualization: J.-Y.K., P.T.T., and J.-Y.P.

Methodology: J.-Y.K., and J.-Y.P.

Investigation: J.-Y.K., S.P., H.C.

Visualization: J.-Y.K., P.T.T., S.P.

Project administration: J.-Y.P.

Supervision: J.-Y.P.

Writing – original draft: J.-Y.K., and J.-Y.P.

Writing – review & editing: J.-Y.K., P.T.T., S.P., H.C., and J.-Y.P.

## Competing interests

The authors declare no competing interests.

## Data and materials availability

The data reported in this work can be provided upon request to corresponding authors.

## Supplementary Materials

Materials and Methods

Figs. S1 to S6

References (*83–88*)

## Notes

### Competing Interest Statement

The authors have declared no competing interest.

### Summary of Updates

Updated missing words in the introduction section.

