## Supplementary Materials for "Direct imaging of neural activity reveals neural circuits via spatiotemporal activation mapping"

**Materials and Methods**

Animals

Six C57BL/6J wild-type mice (29 ± 4 g, 3 – 3.5 month old, Charles River Laboratory, Saint-Germain-Nuelles, France) were used in the experiments. This study was carried out with approval by the Institutional Animal Care and Use Committee (APAFIS#18001-2018121010505627 v1).

Experiments

All experiments for DIANA and BOLD fMRI were performed using a surface cryoprobe on a 11.7 T animal scanner (BioSpec 117/16 USR, Bruker, Ettlingen, Germany). MRI experiments for phantom imaging to test the signal generated by trigger delay were performed on a 9.4 T animal scanner (BioSpec 94/30 USR, Bruker).

*Animal preparation.* Mice were initially anesthetized with 4% isoflurane and maintained at 2% during animal preparation and were placed prone on a heat-regulated cradle. Xylocaine droplets were applied in the ears as a local anesthetic, and ear bars and a tooth bar were used to secure the animal's head still in a nose cone. The mice were self-breathing with a continuous supply of oxygen and air gases (1:1 ratio) through the nose cone at a rate of 1 liter/min. A subcutaneous catheter was placed in the back to deliver a 0.3 mg/kg bolus of medetomidine (Domitor®, Vetoquinol) followed by a continuous infusion at 0.6 mg/kg/h. The isoflurane level was slowly reduced over 5 min until complete discontinuation. Rectal temperature (37 °C) and respiration rate (140 – 160 bpm) were controlled with the SAII's MR-compatible Model 1030 Small Animal Monitoring and Gating System (SA Instrument, Inc., Stony Brook, NY, USA).

*BOLD fMRI.* Two-dimensional (2D) gradient-echo echo planar imaging (EPI) was used for BOLD fMRI. Scan parameters were as follows: repetition time/echo time (TR/TE), 1000/15 ms; flip angle (FA), 45°; field of view (FOV), 24 × 12 $mm^2$; matrix size, 128 × 64; dummy scans, 40 TR; and slice thickness, 0.5 mm or 0.8 mm. First, to check the location of sensory thalamic areas and forelimb primary somatosensory cortex (S1FL), ten 0.5 mm thick coronal slices including the thalamus and S1Fl were acquired. Electrical stimulation of 0.5 mA current strength, 2 ms pulse duration, and 10 Hz frequency was delivered to the right forepaw during the scan, using stainless steel electrodes connected to the isolated pulse stimulator (A-M System Model 2100). The stimulation paradigm consisted of 5 blocks of 20s-on/20s-off cycles. When examining which coronal slices the thalamus and S1FL responded to, BOLD responses in the S1FL and thalamus were approximately 4 to 5 coronal slices apart. Based on these results, a single 0.8 mm thick oblique slice was set using the same stimulation paradigm to observe BOLD responses in the thalamus and S1FL simultaneously. The oblique slice was acquired before and after BOLD fMRI experiments to determine whether or not the mouse responded to the stimulation during the DIANA fMRI experiment. A total of 3 runs were repeated.

*DIANA fMRI.* Experiments were performed using 2D fast low-angle shot (FLASH) based line-scan imaging sequence (ParaVision 6.0.1). Scan parameters were as follows: TR/TE = 5/2 ms; FA, 4°; FOV, 25.6 × 12.8 $mm^2$; matrix size, 128 × 64; slice thickness, 0.8 mm; bandwidth, 75kHz; and scan time, 12.8 s/trial. Sufficient dummy scans (8s, 1600 TR) were used to achieve steady-state magnetization prior to main sequence. Both gradient and RF spoiling were used to suppress the effects of residual transverse magnetization. RF spoiling was applied by default module in Bruker Paravision 6.0.1 software, that is, using a quadratic phase cycling schedule to increment the phase of $j^{\text{th}}$ excitation RF pulse by $j \times 117°$. Therefore, the phase of each excitation RF pulse was given by

$$\Phi_j = \frac{1}{2}\Phi_0(j^2 + j + 2), \quad j = 0, 1, 2, \cdots, \qquad [1]$$

where $\Phi_0 = 117°$. In this case, the phase of the RF pulse increases at each k-space line of the time series within the interstimulus interval, whereas for conventional RF spoiling it increases at each phase encoding step. An identical 0.8 mm thick oblique slice acquired in BOLD fMRI experiment was acquired for DIANA fMRI with 0.5 mA current strength, 1 ms pulse duration, and 5 Hz frequency. The stimulus was repeated 64 times, corresponding to the number of phase-encoding steps with an interstimulus interval of 200ms. The stimulation paradigm consisted of 50 ms pre-stimulation, 1 ms stimulation, and 149 ms post-stimulation. 40 trials per mouse were acquired and used for analysis. To minimize neural adaptation, a 120-second rest period was provided every five trials.

*Phantom imaging.* A volume coil with an inner diameter of 86 mm and a surface coil with a diameter of 10 mm were used for RF transmission and signal reception, respectively. An identical line-scan imaging sequence used in DIANA fMRI experiments was used for phantom imaging with the same scan parameters except for dummy scans. A water phantom and another water phantom containing trace amounts of gadolinium were prepared and scanned with and without RF spoiling. Gradient spoiling was also applied in both conditions. To achieve steady-state magnetization, more dummy scans (2000 TR, 10 s) were performed considering the longer $T_1$ relaxation time of the water phantom compared to the mouse brain. 40 trials were acquired for each condition and each phantom.

*$T_2$ calculation.* Multi spin-echo (MSE) sequence with 5 TEs was used for $T_2$ calculation. Scan parameters were as follows: TR, 1500 ms; TE, 12/24/36/48/60 ms; FOV, 20 × 20 mm$^2$; matrix size, 128 × 128; slice thickness, 3 mm. $T_2$ was calculated using simple mono-exponential decay fitting based on Equation [1].

$$S(n) = S_0 e^{-n\frac{TE}{T2}}, \; n = 1, 2, \cdots, 5, \qquad [1]$$

where $n$ is the number of echoes. Due to the effect of stimulated echo (83), the first echo was excluded for mono-exponential decay fitting (84).

Data analysis

All DIANA and BOLD fMRI data were processed using home-built MATLAB (R2022b, MathWorks), Analysis of Functional Neuroimages package (AFNI) (*85*), and FMRIB Software Library (FSL) (*86*). All analyses of DIANA and BOLD fMRI data were referenced to the Allen Mouse Brain Atlas (*29*) (Allen Brain Institute, http://mouse.brain-map.org).

*BOLD-fMRI.* Individual BOLD activation maps were generated using preprocessing and a general linear model (GLM) analysis with 5 boxcar blocks of 20s-on/20s-off cycles after averaging

3 runs. Preprocessing steps were as follows: linear detrending to remove signal drift and time course normalization relative to the first image. After the preprocessing and GLM analysis, individual BOLD activation maps were generated with a statistical threshold of uncorrected $p < 0.05$ and cluster size > 5 and overlaid on the EPI images. Additional spatial smoothing was not used to generate BOLD activation maps. Time courses were extracted from circular ROIs of ~0.6 mm (3.2 pixels) radius in the thalamus and S1FL regions.

*DIANA fMRI.* For group analysis, each DIANA image of mice #2 - 6 (except for mouse #5) was linearly registered to the DIANA image of mouse #1 using FMRIB's Linear Image Registration Tool (FLIRT) function in FSL. Mouse #5 data were excluded from this analysis because mouse #5 did not show BOLD activation in the thalamus. Transformation matrix for each mouse was extracted by linearly registering the DIANA images averaged across all 40 frames and 40 trials for each mouse to the DIANA image of mouse #1 averaged across all 40 frames and 40 trials, using tri-linear interpolation. Next, each transformation matrix was applied to all 40 frames and 40 trials for corresponding mouse using 'applyxfm' function in FSL.

Spatiotemporal activation mapping was performed to obtain DIANA-activated specific brain regions. Prior to spatiotemporal activation mapping, several preprocessing steps were taken: Voxel-wise temporal smoothing was first applied using a three-point Gaussian kernel, and each frame was spatially smoothed with a Gaussian kernel of 0.33 mm full-width-half-maximum (FWHM) instead of ROI averaging. After that, time series in each voxel was normalized relative to the mean of the pre-stimulation period, and linear detrending was applied to the time series to remove signal drift. Finally, voxels with at least one data point with z-score > 3 or < -3 were excluded as outliers, considering the histogram of all data points in the time series of all voxels. These outlier voxels were highly concentrated in the lower part of the brain, where the temporal signal-to-noise ratio (tSNR) is relatively low because a surface coil was placed on top of the brain. After the preprocessing, a peak response time map was generated by calculating the time that has maximum amplitude of the time series in each voxel within the brain. Next, spatiotemporal clustering was performed by clustering the voxels with the same peak response time and selecting clusters with size greater than 4 for each time frame. Then, to test the statistical significance of the DIANA response in each cluster, statistical comparisons were made between the baseline mean of the pre-stimulus period and every post-stimulus time point in the average time series for each cluster using data from all five rats. Through this statistical process, clusters with at least one statistically significant data point survived to form the spatiotemporal activation map. Using the spatiotemporal activation map, activation voxels as clusters were defined in the forelimb sensory processing regions of interest. Finally, time series were extracted from those defined ROIs using temporal smoothing using a three-point Gaussian kernel, region-of-interest (ROI) averaging instead of spatial smoothing, signal normalization, and linear detrending. The DIANA response latency for each mouse was calculated for the first peak above the mean + 0.7 × standard deviation (SD) within a 45 ms window size, from 20 ms before to 25 ms after the time of the maximum peak of the average time series. Additionally, the control ROI was defined within the high tSNR muscle region with size 3 × 3.

For the heatmaps, after interpolating the data to 10 times higher spatial resolution using linear interpolation, rectangular ROIs containing activation areas in each of thalamic nuclei (VPL and POm) and cortical (S1 and S2) regions were masked, respectively, and rotated at an appropriate angle to be vertical. Next, the same preprocessing used for time series analysis was also performed to generate the heatmaps. Additionally, to better visualize the DIANA activations, a 2D median filter with kernel size of 13 × 13 voxels was applied to the masked region.

*Simulation.* To test our hypothesis that the DIANA signal has pseudo-random components that are more effectively suppressed by cross-subject averaging, we first assumed that these pseudo-random components are oscillatory signals with a specific frequency and phase for each subject. Then, we simulated artificial signals consisting of deterministic neuronal responses, non-random oscillations of a specific frequency and phase range, and random noise. For deterministic neuronal responses, identical 40 time series with 2-point peak responses were generated for each mouse, and peak amplitudes in each trial were multiplied by an exponential decay function to mathematically reflect neuronal adaptation (*69*). The peak response $R(n)$ was given by

$$R(n) = \left(a * e^{-\frac{f}{f_c}} + (1 - a)\right)^{n-1}, \; n = 1, 2, \cdots, 40, \qquad [2]$$

where $n$ is the trial number. The parameters were as follows: $a$ = 0.585, $f_c$ = 8.663 Hz, and $f$ = 5 Hz. For the nonrandom oscillatory component, 40 sinusoidal signals were generated, with amplitudes randomly chosen within (0, 1] for each mouse, frequencies randomly chosen between 0.5 and 3.5 Hz for each mouse assuming slow-wave activity under anesthesia (*87*), phases (φ) randomly chosen within [0, 2π] for each mouse, and phases randomly chosen within [φ, φ+π] for each trial of each mouse. Finally, random noises were generated from a Gaussian distribution with SD 0.33 and mean 0. Among the artificially generated trials, trials with a multiple of 5, that is, 5 to 40 trials, were randomly selected from a randomly selected single mouse, and for comparison, 1 trial to 8 trials (i.e., 5 to 40 trials divided by 5) were randomly selected from each of the 5 mice, resulting in a total of 40. Then, the SD of the pre-stimulation period was averaged over 100,000 iterations. The same calculations were performed with *in vivo* data acquired at 11.7 T.

<u>Statistics</u>

For all statistical comparisons, Shapiro-Wilk test was performed to test normality except for samples not less than 30. The samples with more than 30 were considered to follow a normal distribution based on the central limit theorem (*88*). For all statistical comparisons between the two groups satisfying normality, a Levene's test was performed to check for homoscedasticity, followed by a one-tailed paired *t*-test or Welch's *t*-test. If normality was not satisfied in this case, a one-tailed Wilcoxon signed-rank test was performed. For all statistical comparisons between three group satisfying normality and homoscedasticity were conducted with one-way analysis of variance (ANOVA) with Fisher's LSD *post hoc* test. In cases where normality was not satisfied, Kruskal-Wallis ANOVA with Fisher's LSD *post hoc* test was used for three group comparisons. Quantitative data were all expressed as mean ± standard error of the mean (SEM) or mean ± standard deviation (SD).

### Supplementary figures

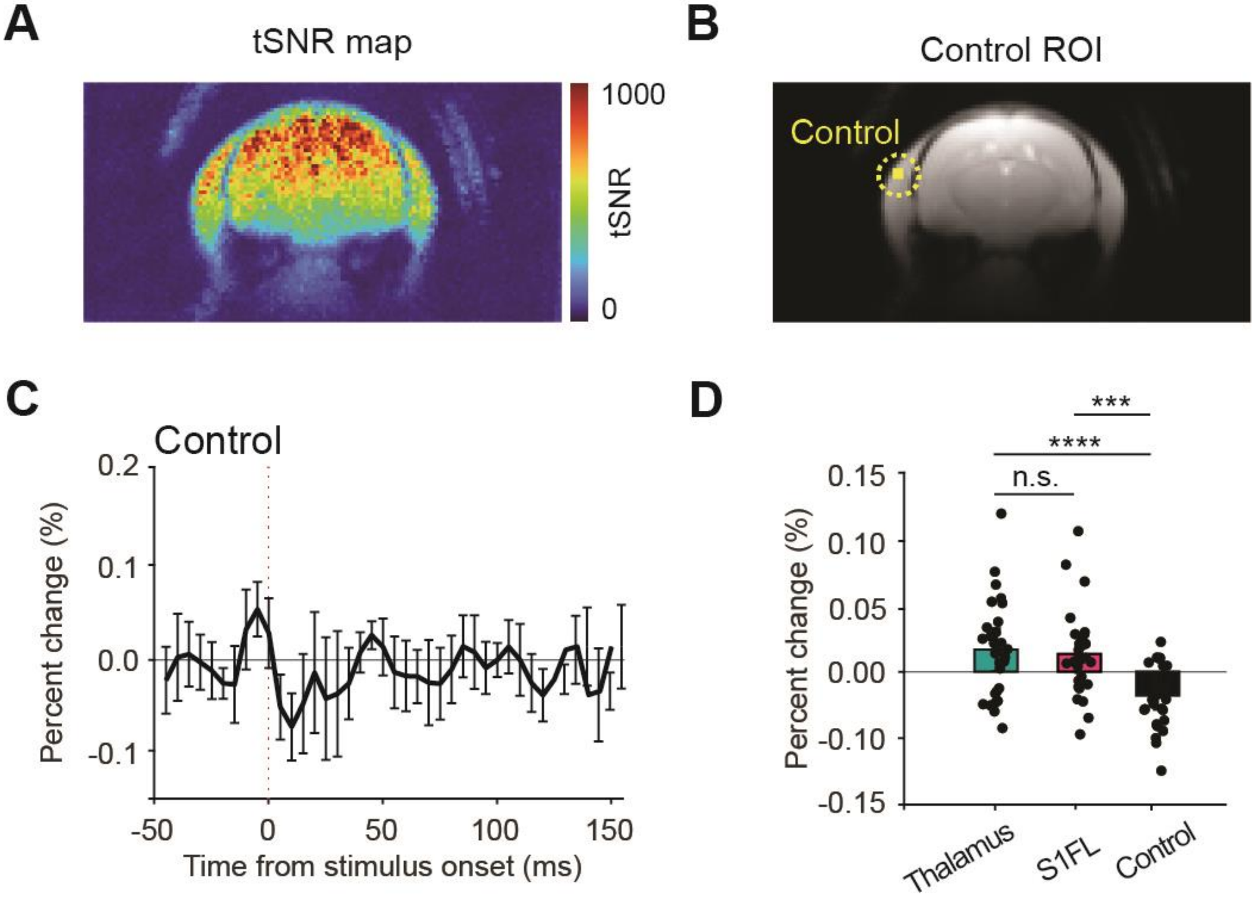


**Fig. S1. tSNR map and DIANA response in control region. (A)** tSNR map of the average DIANA image for 200 trials (= 5 mice × 40 trials/mouse). **(B)** Control ROI in the muscle region with high tSNR. **(C to D)** Time series in the control region (C) and mean percent signal changes during post-stimulation for the thalamus (cyan), S1FL (magenta), and control region (black) (D). The vertical dotted red line indicates the onset time of electrical forelimb stimulation (C). All data are mean ± SEM. ***: $p < 0.001$, ****: $p < 0.0001$, n.s. > 0.05 for one-way ANOVA with Fisher's LSD *post hoc* test.

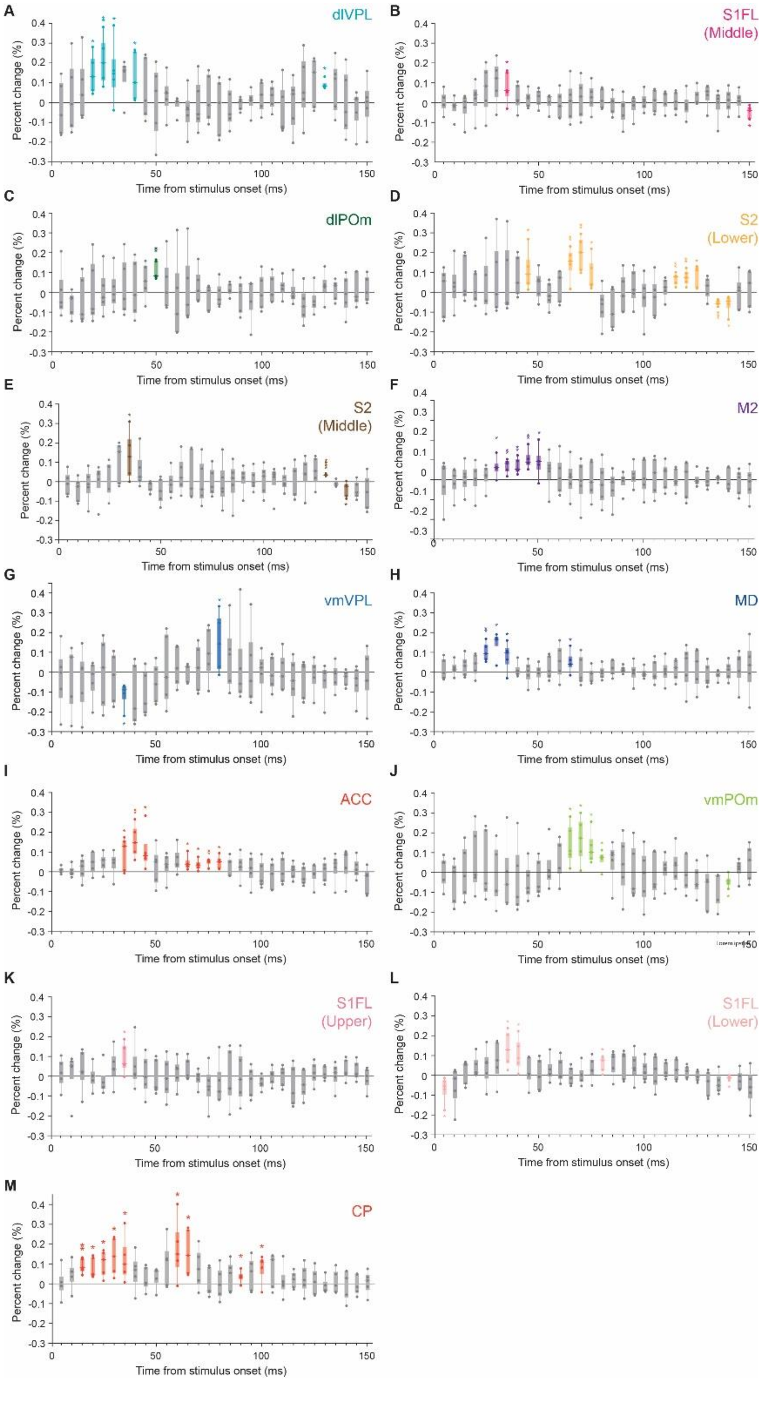


**Fig. S2**. **Statistical significance of DIANA signal changes at each time point. (A to M)** DIANA responses in dorsolateral VPL (dlVPL) (A), middle S1FL (B), dorsolateral POm (dlPOm) (C), lower S2 (D), middle S2 (E), M2 (F), ventromedial VPL (vmVPL) (G), MD (H), ACC (I), ventromedial POm (vmPOm) (J), upper S1FL (K), lower S1FL (L), and CP (M) ($n$ = 5 mice). In the box plots, each box represents 25th to 75th percentiles, the horizontal line represents the median, and whisker represents the range from minimum to maximum. *: $p < 0.05$, **: $p < 0.01$, ***: $p < 0.001$, ****: $p < 0.0001$ for one-tailed Wilcoxon signed-rank test or one-tailed Welch's $t$-test.

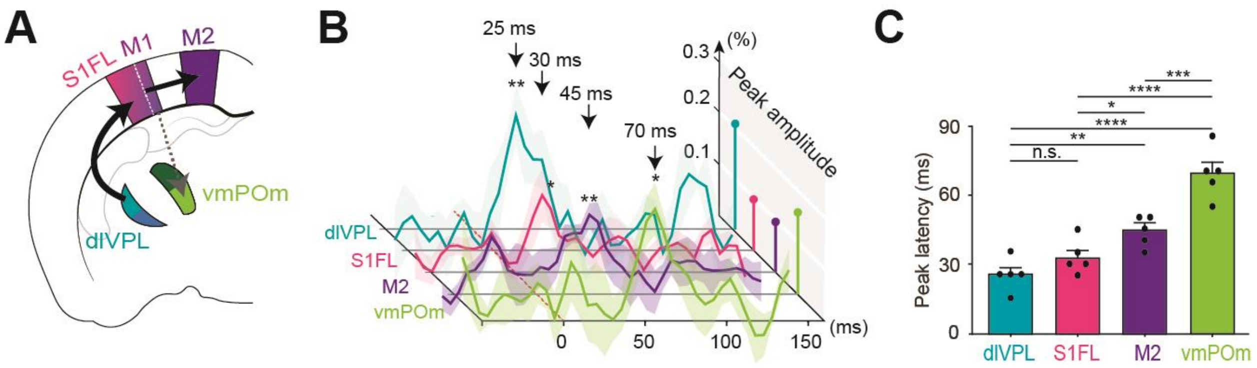


**Fig. S3. Extended Circuit IV including CT feedback projection to POm. (A)** Illustration of TC and CC projections from the dorsolateral VPL (dlVPL) through S1FL and M1 to the M2, and additional CT projection from M1 to the ventromedial POm (vmPOm, light green). **(B)** DIANA time series and **(C)** mean latencies of peak DIANA responses in dlVPL, S1FL, M2, and vmPOm ($n$ = 5 mice). The dotted red line indicates the onset time of electrical forelimb stimulation (B). In time series plots, the maximum DIANA response amplitude for each ROI was displayed as a percentage change in the right-end plane in projection form. All data are mean ± SEM. n.s.: $p > 0.05$, *: $p < 0.05$, **: $p < 0.01$, ***: $p < 0.001$, and ****: $p < 0.0001$ for one-way ANOVA with Fisher's LSD *post hoc* test.

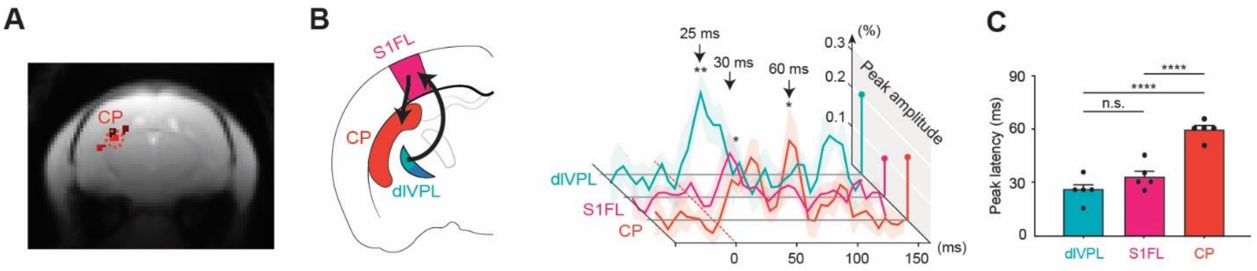


**Fig. S4. Cortico-striatal (CS) pathway involved in forelimb sensory processing. (A)** Four activated regions in caudoputamen (CP) (red). **(B)** Illustration of CS projection from S1FL (magenta) to the CP subregion with the highest response (red) and DIANA time series (right). **(C)** Mean latencies of peak DIANA responses in the dorsolateral VPL (dlVPL), S1FL and CP ($n$ = 5 mice). Dotted red lines indicate the onset time of electrical forelimb stimulation (B). All data are mean ± SEM. n.s.: $p > 0.05$, ****: $p < 0.0001$ for one-way ANOVA with Fisher's LSD *post hoc* test.

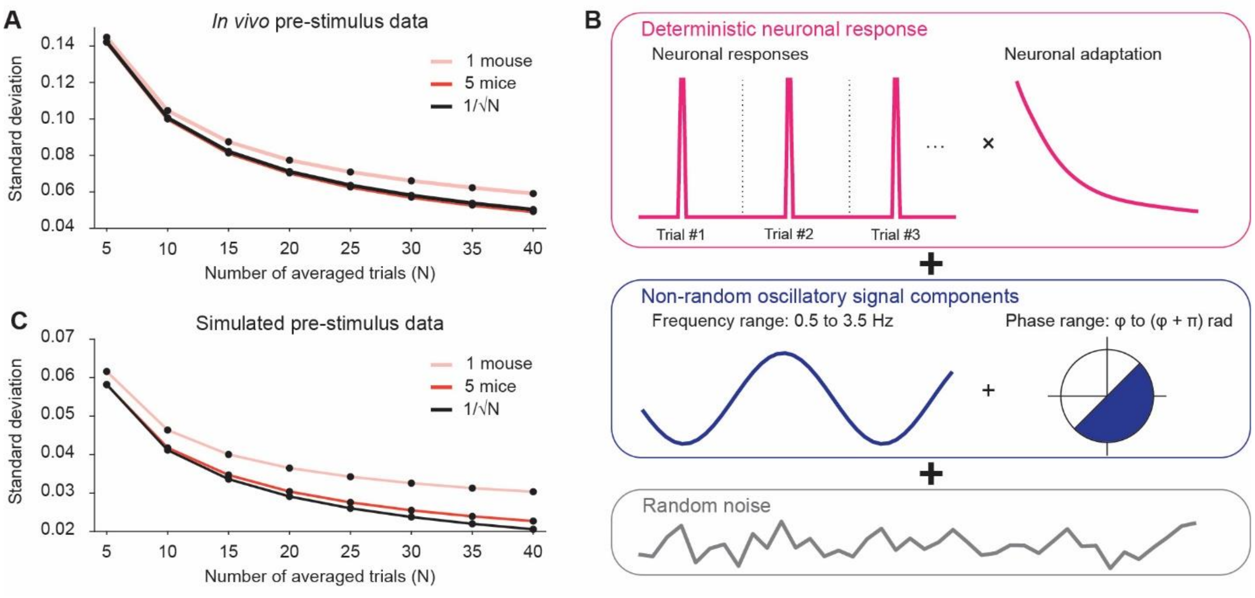


**Fig. S5. Comparison of DIANA signal baseline fluctuation between between-subject and within-subject averaging. (A)** Standard deviation calculated from the pre-stimulus period of the DIANA fMRI time series with respect to the number of averaged trials. **(B)** Artificial signals consisting of deterministic neuronal responses undergoing neuronal adaptation (magenta), non-random oscillatory signals with specific frequency and phase ranges (blue), and random noise (gray). **(C)** Standard deviation calculated from the pre-stimulus period of the artificial signals with respect to the number of averaged trials. The simulated pattern of baseline fluctuations in the averaged data (C) is similar to that in the averaged *in vivo* data (A).

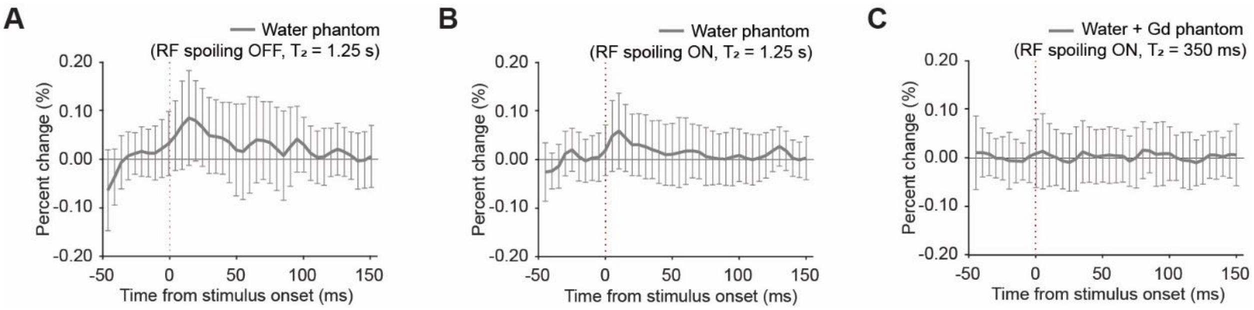


**Fig. S6. Signals produced by trigger delay in water phantom experiments. (A)** DIANA time series obtained using a water phantom ($T_2$ = 1.25 s) when RF spoiling was turned off. The signal change generated by the trigger delay (25 µs in our experiment) was observed to peak at ~15 ms (0.083%). **(B)** DIANA time series obtained using a water phantom when RF spoiling was turned on. The signal change generated by the trigger delay was observed to peak at ~10 ms (0.056%). **(C)** DIANA time series obtained using a Gd-containing water phantom ($T_2$ = 350 ms) when RF spoiling was turned on. Signal changes due to trigger delay were well suppressed (0.00080 ± 0.0066% for all time points). The red vertical dotted line indicates the trigger onset time.
